# Injury-induced cold sensitization in *Drosophila* larvae involves behavioral shifts that require the TRP channels Pkd2 and Brv1

**DOI:** 10.1101/342717

**Authors:** Heather N. Turner, Atit A. Patel, Daniel N. Cox, Michael J. Galko

## Abstract

Nociceptive sensitization involves an increase in responsiveness of pain sensing neurons to sensory stimuli, typically through the lowering of their nociceptive threshold. Nociceptive sensitization is common following tissue damage, inflammation, and disease and serves to protect the affected area while it heals. Organisms can become sensitized to a range of noxious and innocuous stimuli, including thermal stimuli. The basic mechanisms underlying sensitization to warm or painfully hot stimuli have begun to be elucidated, however, sensitization to cold is not well understood. Here, we develop a *Drosophila* assay to study cold sensitization after UV-induced epidermal damage in larvae. Larvae respond to acute cold stimuli with a set of unique behaviors that include a contraction of the head and tail (CT) or a raising of the head and tail into a U-Shape (US). Under baseline, non-injured conditions larvae primarily produce a CT response to an acute cold (10 °C) stimulus, however, we show that cold-evoked responses shift following tissue damage: CT responses decrease, US responses increase and some larvae exhibit a lateral body roll (BR) that is typically only observed in response to high temperature and noxious mechanical stimuli. At the cellular level, class III neurons are required for the decrease in CT, chordotonal neurons are required for the increase in US, and chordotonal and class IV neurons are required for the appearance of BR responses after UV. At the molecular level, we found that the transient receptor potential (TRP) channels Polycystic kidney disease gene 2 (*Pkd2*) and *brivido-1* (*brv1*) are required for these behavioral shifts. Our *Drosophila* model will enable a sophisticated molecular genetic dissection of genes and circuits involved in cold nociceptive sensitization.

## Introduction

Nociceptive sensitization is an exaggerated behavioral or biological response to a normal stimulus due to a lowered nociceptive threshold. It is typically observed after tissue damage or injury. Nociceptive sensitization commonly develops near the site of injury, where the local sensory neurons are temporarily hypersensitized until the wound effectively heals [1]. Nociceptive sensitization is thought to foster protective behavioral mechanisms to prevent further tissue damage and aid the healing process [2]. However, when the pain extends beyond the time necessary for wound healing, it becomes maladaptive and more difficult to treat. There are a number of clinical conditions that are associated with maladaptive nociceptive sensitization, which greatly reduce the quality of life in patients and remain difficult to treat. Thermal sensitivity to cool-cold temperatures can be experienced as innocuous cool temperatures being perceived as painful (cold allodynia), or harsh cold temperatures perceived as *more* painful (cold hyperalgesia). While cold allodynia is common in patients with multiple sclerosis [3], fibromyalgia [4], stroke [5,6], and chemotherapy-induced neuropathy [7,8], the mechanisms that underlie cold sensitization in these conditions are largely unknown.

A wide array of nociceptive assays and tools have been used in vertebrates to investigate cold nociception and nociceptive sensitization, including: a cold plate [9], tail-flick [10], exposure to acetone [11], or dry ice assays [12]. Under non-injured conditions vertebrate cool-sensing neurons (peripheral C and A*δ* fibers) are activated between 20-37 °C with increased firing upon cooling down to 20-17 °C, while noxious cold-sensing neurons are activated between 10-20 °C with increased firing down to 0 °C [13]. These neurons are also required for observed sensitized responses to cold [13], however neuronal activation and hence nociceptive thresholds are prone to shift under the context of injury or damage [14] and the mechanisms underlying these shifts are largely unknown.

At the molecular level, a handful of cold-sensitive channels have been proposed as cool or noxious cold receptors. A large focus has been on transient receptor potential (TRP) channels, which have known functions in nociception, thermosensation, and nociceptive sensitization [15,16]. TRP channels are variably selective cation channels containing multiple subunits and six transmembrane domains that fall into several different gene families including: TRPC, TRPM, TRPV, TRPA, TRPP and TRPML. While TRPV channels have been characterized as warmth and heat activated [17], TRPM8 and TRPA1 are most notably implicated in cold sensing [18,19] and inflammatory or damage-induced cold hypersensitivity in vertebrates [20,21]. Experiments in TRPM8 null mice, however, have shown that a small population of dorsal root ganglion (DRG) neurons lacking TRPM8 are still able to respond to cold or menthol [22]. Likewise mice mutant for TRPM8, while exhibiting significant impairment of cold sensing, will still avoid painfully cold temperatures (16-20 °C) [22]. Similar studies in TRPA1 null mice show that although these mice show severe defects in responding to cold, the mutation does not completely abolish cold-evoked behaviors [23]. Lastly, additional studies of cold-responsive DRG (as well as superior cervical ganglia) neurons did not respond to TRPM8 or TRPA1 agonists [24]. Together these studies suggest that cold receptor(s) beyond TRPM8 and TRPA1 exist.

Studies in noxious cold detection and cold hypersensitivity in vertebrates to date suggest a complex process, likely involving multiple receptors and/or modulators. Genetically tractable organisms such as the fruit fly could greatly aid this work by offering a simpler platform for study that offers a sophisticated genetic toolkit. Identifying the cells and channels involved in cold-sensing in invertebrates is relatively recent, including studies in flies [25,26], worms [25,27], and the leech [28]. The majority of these studies (other than [26]) however, focused on fairly innocuous ‘cool’ ranges (> 12 °C) and measured thermal preference, or avoidance of temperatures just outside preferred ranges. *Drosophila* utilize cool-sensing neurons in the head [29,30], and chordotonal neurons in the larval body wall [31], as well as the TRP channels *trp*, *trpl* [32], *inactive* [31], and the TRPP type *brivido* genes [30], to avoid mildly cool temperatures outside their preferred temperature range.

Recently, it was shown that *Drosophila* larvae also respond to acute noxious cold stimuli with distinct behaviors, including a contraction (CT) of the anterior and posterior segments towards the center of the body, and a raising of these anterior and posterior segments into the air to create a U-Shape (US) [26]. Both of these behaviors are distinct from normal locomotion, most gentle touch behaviors [33-36] as well as the stereotyped 360° lateral body roll (BR) behavior observed in response to high heat [37,38] and noxious mechanical stimuli [37,39-41]. The primary cold-evoked behavior, CT, requires class III multidendritic (md) sensory neurons, which innervate the barrier epidermis and are directly activated by cold temperatures [26]. Cold-evoked CT behavior is also mediated by class III expression of the TRP channels Polycystic kidney disease gene 2 (Pkd2, a TRPP channel), NompC (a TRPN channel) and Trpm [26].

Although invertebrate models including *C. elegans* [42], *Aplysia* [43], and *Drosophila* [38,44,45] have been used to study nociceptive sensitization, none have looked at injury-induced cold hypersensitization. In *Drosophila,* larvae sensitize to mildly warm and hot stimuli following UV-induced tissue damage [38,45]. Larvae normally respond to acute high temperature stimuli (≥45 °C) with a 360° lateral BR behavior [38]. After UV damage however, larvae exhibit BR in response to subthreshold temperatures (thermal allodynia), and show a robust increase in the number of BR responders to noxious heat with a concomitant decrease in response latency (thermal hyperalgesia) [38]. Sensitization to warm and hot stimuli require class IV md sensory neurons, which are also required for baseline responses to high temperature in the absence of tissue damage [38]. UV exposure results in a rapid apoptotic breakdown of the larval epidermis between 16-24 hours after UV administration; the underlying sensory neurons, however, remain intact [38,46]. Further, class IV md neurons expressing the TRP channel *painless* (a TRPA channel) are required for baseline responses and UV-induced sensitized responses to heat [45].

Currently, it is unknown if *Drosophila* larvae sensitize to cold stimuli following tissue damage, and if so, whether they utilize known or distinct sensory neurons and receptors for this sensitization. Here we characterize a marked shift in behavioral responses to cold temperatures after epidermal injury. We have found that class III and class IV md sensory neurons, as well as peripheral chordotonal neurons, are required for this observed shift in cold-evoked behavior. Lastly, we identify a role for the TRP channels Pkd2 and Brv1 in injury-induced shifts in behavioral responses to cold temperatures. This work establishes a platform for future studies on the cellular and molecular bases of cold nociception and injury-induced cold sensitization.

## Materials and Methods

### Fly Stocks and Genetics

*Drosophila melanogaster* larvae were raised on cornmeal media at 25°C. *w^1118^* was used as a control strain. Mutants: *Pkd2^1^*[47] and *brv1^L563>STOP^* [30] (a gift from Marco Gallio). Deficiencies *Df(2L)BSC407* (*Pkd2*) from Bloomington *Drosophila* Stock Center and *brv1-Df* (*Df(3L)Exel9007*) from Marco Gallio. Gal4 lines: *19-12-Gal4* (class III[48]), *ppk1.9*-*Gal4* (class IV [49]), and *iav-Gal4* (chordotonal [50,51]). UAS transgenes: *UAS-TeTxLC* (active tetanus toxin [52]), *UAS-IMP TNT VI-A* (inactive tetanus toxin [52]), *UAS-mCD8::GFP* [53] and *UAS-GCaMP6m* [54]. *UAS-RNAi* line from Vienna *Drosophila* RNAi Center [51]: 6940 (*UAS-Pkd2^RNAi^*; from Bloomington *Drosophila* Stock Center [55]: 31496 (*UAS-brv1^RNAi^*).

### Local cold probe assay

Details on the custom-built Peltier cold probe (TE Technologies, Inc.) used in this study can be found here [26]. For behavioral analysis, mid 3^rd^ instar larvae were placed on a stage under a brightfield stereomicroscope (Zeiss Stemi 2000). As previously described [26], the probe tip was gently placed on the dorsal midline (segments A4-A6) for up to 10 s or the initiation of a behavioral response, while larvae moved freely under the microscope. Larvae that did not exhibit a behavioral response within 10 s were recorded as non-responders. Behavioral responses were characterized as either: 1. A contraction (CT); 2. A 45-90 ° simultaneous head and posterior raise (US for U-shape) (as described previously [26]); or 3. A 360 ° lateral body roll (BR), which was observed under sensitized conditions. For all data, only larvae that initially exhibited normal locomotion were tested and each larva was stimulated only once. Lastly, in *Gal4/UAS* experiments, transgenes were heterozygous and no balancers or markers were present in the larvae used for behavioral testing. Experimenter was blind to genoype being tested.

### UV-induced Tissue Damage

To determine effects of UV-induced tissue damage on larval cold nociception and sensitization, larvae were irradiated by exposure to 10-14 mJ/cm^2^ UV-C (as previously described [38]) and then allowed to recover on food in a 25 °C incubator before being tested in the nociceptive assays 4, 8, 16 or 24 hours later. For this, 3^rd^ instar larvae of similar size were selected 4-5 days after egg laying and irradiated, then tested in the cold assay after recovery from UV. Before irradiation, larvae were immobilized on a cold slide for a few minutes, then fine-tipped forceps were used to position the larvae dorsal side up in a row along the length of the slide. The spectrolinker XL-1000 UV crosslinker (Spectroline) was warmed up, and the UV dose was measured with a hand-held UV spectrophotometer (AccuMAX XS-254, Spectroline) just prior to exposure to get an accurate reading. All UV doses fell within 10-14 mJ/cm^2^, which have been shown to induce epidermal cell death [56] while sparing the peripheral sensory neurons from significant morphological changes [46,57]. Larvae were then administered UV (actual dose recorded), and carefully placed into a vial of food with a paintbrush to recover for a variable amount of time before being tested in nociceptive assays.

### Neuronal morphology: live imaging and confocal microscopy

Live confocal imaging of neuronal morphology was performed as previously described [26,58]. Briefly, third instar larvae were mounted on microscope slides using 1:5 (v/v) diethyl ether:halocarbon oil and imaged using Zeiss LSM780 laser confocal microscope. For neurometric analyses of class III md neurons, maximum intensity projections of z-stacks were analyzed using ImageJ as previously described [59]. For chordotonal neurons, maximum projections were analyzed using Zeiss Zen Blue software, where mean fluorescence intensity (total fluorescence intensity normalized to area) was analyzed for regions of interest (cell body and dendrites). In this study mean fluorescence intensity was used as a measure of potentially altered cellular integrity or potential changes in protein translation, trafficking or recycling that could impact fluorescence intensity.

### *in vivo* calcium imaging

For *in vivo* GCaMP analysis to visualize CIII, CIV, and Ch neurons, *19-12-Gal4* (Class III) and *UAS-GCaMP6m* were used in combination with *ppk1.9-Gal4* (Class IV) or *iav-Gal4* (Chordotonal) respectively. *in vivo* calcium imaging was performed as previously described [26,60]. Briefly, third instar larvae were mounted on a microscope slide with minimal water to prevent desiccation and placed on a Peltier stage (Linkam PE120) for time lapse imaging. The following temperature regimen was used during time lapse imaging: 1 minute at 25°C, ramp down to 6°C at 20°C/minute, hold at 6°C for 10 seconds, ramp up to 25°C at 20°C/minute, and hold at 25°C for one minute. Images were recorded at 212.55 μm × 212.55 μm resolution and 307.2 ms per frame. Raw time-lapse files were motion corrected in Fiji using the Stack Reg function. A region of interest was manually drawn around the cell body and mean fluorescence intensity across time was collected. ΔF/F_0_ was calculated as previously described [26].

## Results

### *Drosophila* larvae are sensitized to cold after tissue damage

Given that *Drosophila* larvae sensitize to noxious heat after tissue damage [38], and respond to noxious cold in the absence of injury [26], we wanted to determine if larvae also sensitize to noxious cold after tissue damage. We used UV irradiation to damage the dorsal epidermis [38] and then allowed mock-treated or irradiated larvae to recover for different amounts of time before testing for changes in cold nociception. Intriguingly, the primary cold behavior at 10 °C, contraction (CT) [25], was significantly decreased at 16 and 24 hours after UV damage compared to mock-treated controls (Fig 1A, S1A and B Fig respectively). Interestingly, the decrease in CT response comes primarily from the number of fast responders (within 3 seconds) rather than slow responders (between 4-10 seconds) at both 16 (S1A Fig) and 24 hrs (S1B Fig) after UV. Despite the observed decrease in CT behavior, upon analyzing cold nociceptive class III (CIII) sensory neurons after mock or UV irradiation, we found they exhibit normal dendritic morphology after UV irradiation (Fig 1C-G). In contrast to the CT response, the percent of U-Shape (US) responders was increased 16 and 24 hours after UV (Fig 1B, S1A and B Fig respectively). Once again, the change in the response was derived primarily from fast responders (S1A & B Fig). Interestingly, we also observed a significant number of larvae that responded to the cold probe with a nocifensive body roll (BR) at these time points (Fig 1B, S1A & B Fig). BR is normally only seen in response to high temperature and harsh touch and is not observed in response to a cold stimulus under baseline conditions [26]. These data suggest that epidermal tissue damage results in a shift of behavioral responses to acute noxious cold stimuli despite the lack of damage to the underlying sensory neurons. In this case, this shift appears as a switch from the dominant cold-responsive behavior at 10 °C, a CT, to US and the normally heat-evoked BR response.

**Fig 1.**
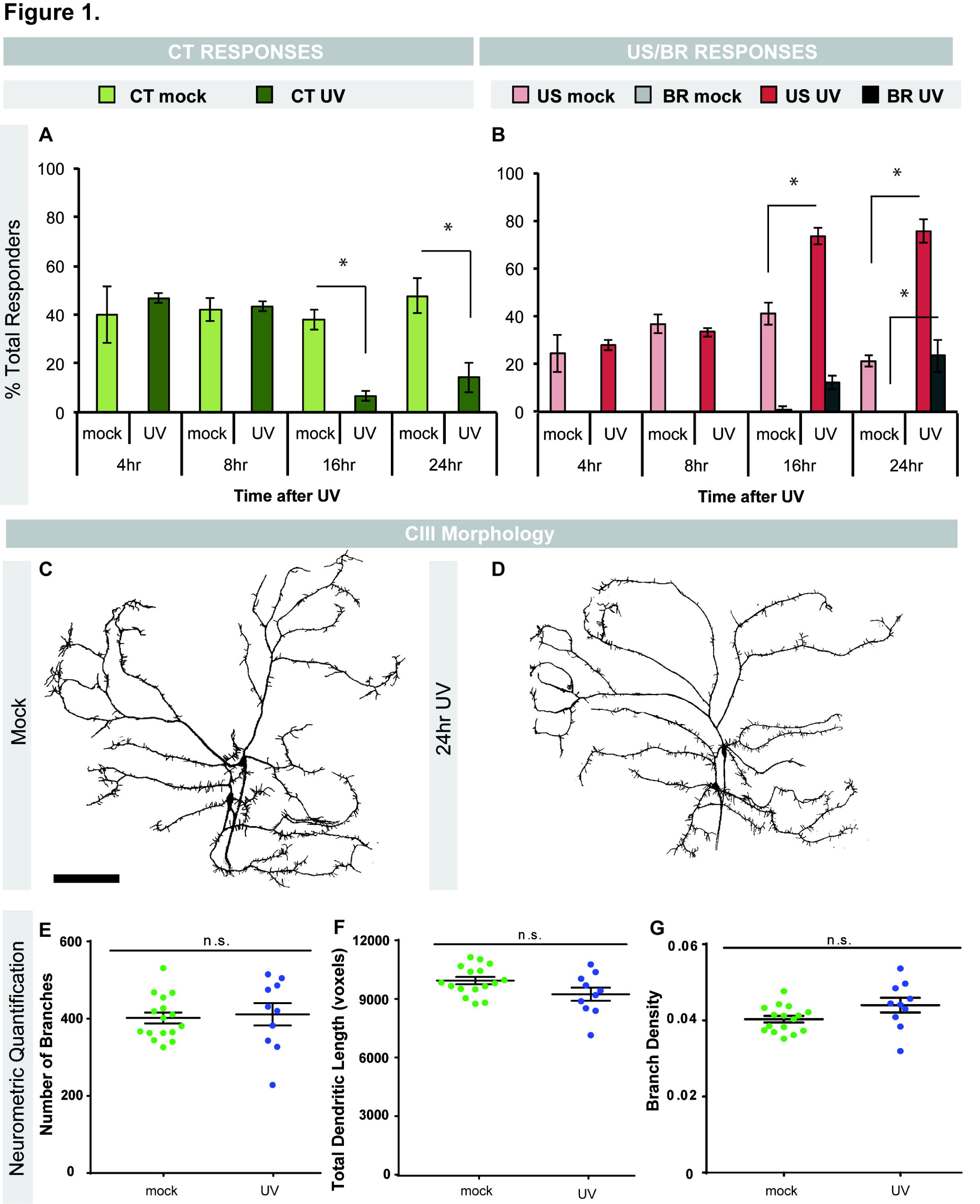
Cold-evoked behavioral switches and neuronal morphology after UV. (A) Percent of CT responders or (B) percent of US and BR responders to cold probe (10 °C stimulation) at indicated times after UV irradiation in mock- or UV-treated larvae. CT = Contraction; US = U-Shape; BR = Body roll, n = 90. (C-D) *in vivo* confocal images of larval CIII sensory neuron dendrite morphology in mock treated larvae (C) and larvae 24 hours post UV treatment (D). Neurons were visualized via *19-12-Gal4>UASmCD8::GFP*. Scale bar: 100 μm (E-G) Neurometric quantification of CIII sensory neurons. (E) number of branches, (F) total dendritic length, and (G) branch density (number of branches/total dendritic length). n = 10-16 neurons. Data are presented as mean ± s.e.m (A-B, E-G). Stats: (A-B) Two-tailed Fisher’s Exact test, * = p < 0.05, comparisons were made between UV and mock control at each time point. (E-G) Two-tailed t-test, n.s. = not significant, comparisons were between UV and mock control of each neurometric measure.

To determine the temperature range(s) over which UV-induced changes in behavioral responses to cold are observed, we tested larvae 24 hours after UV (peak sensitization response) at multiple cold to cool temperatures. While decreases in CT were observed at all temperatures (Fig 2A), increases in US were not observed at 5 °C or 15 °C (Fig 2B), and the emergence of BR was observed only at 10 °C (Fig 2C). 10 °C was the only temperature where all three changes were observed: decrease in CT, increase in US, and emergence of BR (Fig 2). No changes in cold-evoked behaviors were seen with a RT probe, eliminating gentle touch as a contributor to the observed responses (Fig 2).

**Fig 2.**
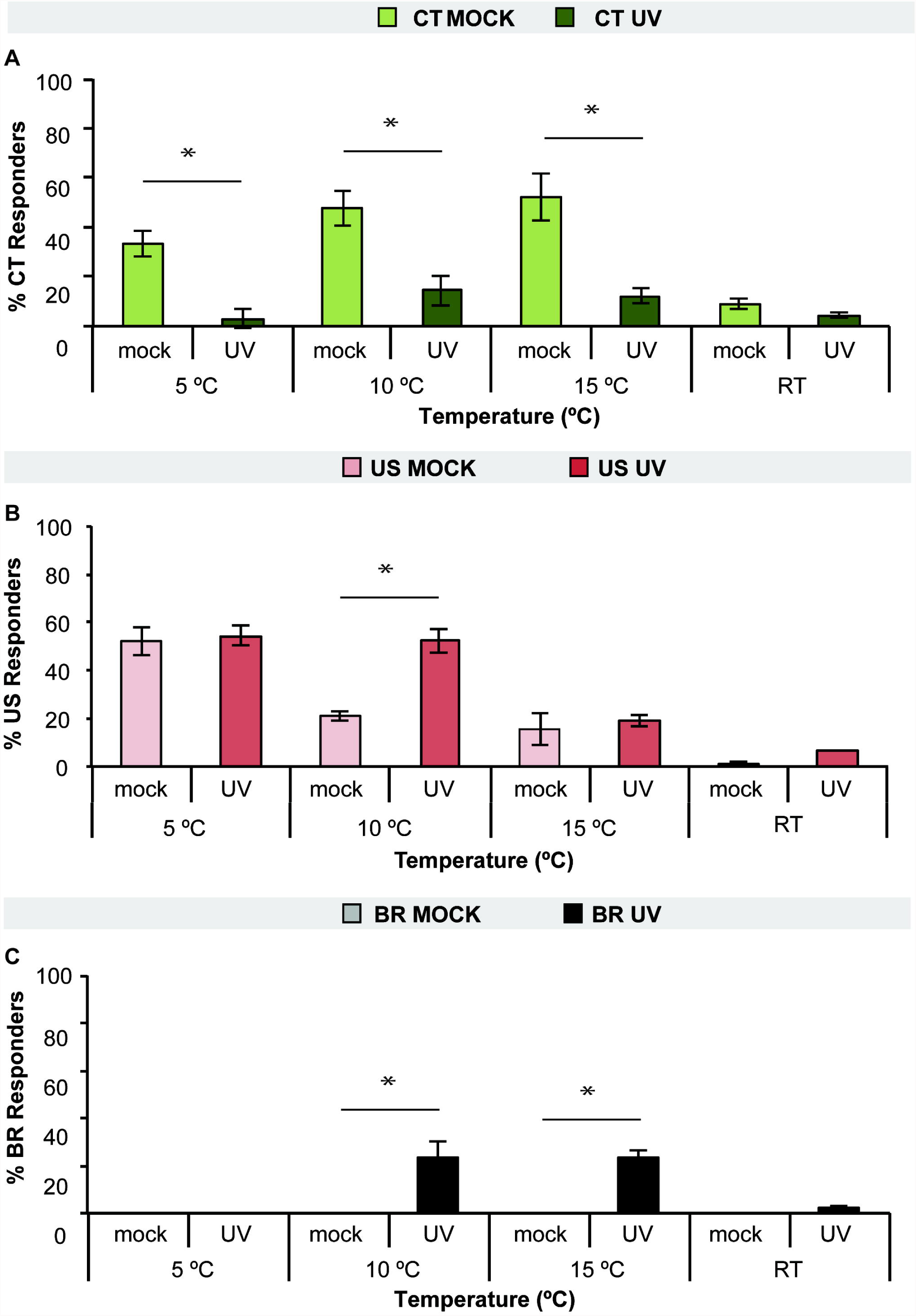
Dependence of UV-Induced cold-evoked behaviors on temperature. Larvae were tested 24 hours after UV exposure or mock treatment with the cold probe either at 5 °C, 10 °C, 15 °C or room temperature (RT) and the percent responders for CT (A), US (B), or BR (C), were averaged. n = 90. Data are presented as the average ± s.e.m.. Stats: two-tailed Fisher’s Exact test, * = p < 0.05, comparisons were made between UV and mock control at each temperature.

Since the received UV dose can vary slightly with each administration (10-14 mJ/cm^2^), we wanted to determine if variations in the UV intensity could impact cold sensitization after UV. Larvae were immobilized (see methods) and subjected to 10-14 mJ/cm^2^ UV to the dorsal side. Larvae were grouped based on the actual measured dose of UV and then allowed to recover for 24 hours before their cold responses were assessed. We observed some changes in CT and BR responses with 13 mJ/cm^2^ UV, however the majority of responses did not differ significantly between doses (S2 Fig). Although the UV dose required to cause apoptosis of the larval epidermis is greater than 12 mJ/cm^2^ [56], our data suggests the degree of apoptosis in the epidermis is not associated with cold-evoked responses after UV.

### Peripheral sensory neurons are required for UV-induced change in cold responses

Since baseline CT responses to cold require CIII md sensory neurons [26] and BR responses to heat and mechanical stimuli are mediated by class IV (CIV) md neurons [37,38], we next determined if either of these neuron classes are required for UV-induced changes in cold responses. We also tested bipolar chordotonal (Ch) neurons which were initially described as stretch receptors [61], but recently found to be important for avoiding cool temperatures in larvae [29,31]. We utilized the *Drosophila* Gal4/UAS system [62] to target the expression of a tetanus toxin transgene (*UAS-TNTE* [52]) in specific classes of sensory neurons using class-specific drivers (see materials and methods). The expression of tetanus toxin essentially prevents neurotransmission, effectively silencing the neurons of interest [52]. We UV-irradiated larvae in which CIII, CIV or Ch neurons were silenced and tested their cold-evoked responses 24 hours after UV exposure. Silencing CIII neurons blocked CT responses in both mock (as shown previously [26]) and UV-treated larvae (Fig 3A). US responses were blocked when Ch neurons were silenced, but not when CIII or CIV neurons were silenced compared to genetic and mock-treated controls (Fig 3B). Consistent with previous studies, increased BR responses after UV were blocked when CIV and Ch neurons were silenced (Fig 3C and [39]). Exposure to UV does not alter CIV neuron morphology [46]. Likewise, analysis of Ch neuron morphology after UV irradiation revealed that were no significant differences in mean fluorescence intensity normalized to area for the cell body or the dendrite (Fig 3D-F). These data suggest a novel and important role for CIV (BR) and Ch neurons (US, BR) in UV-induced changes in cold responses.

**Fig 3.**
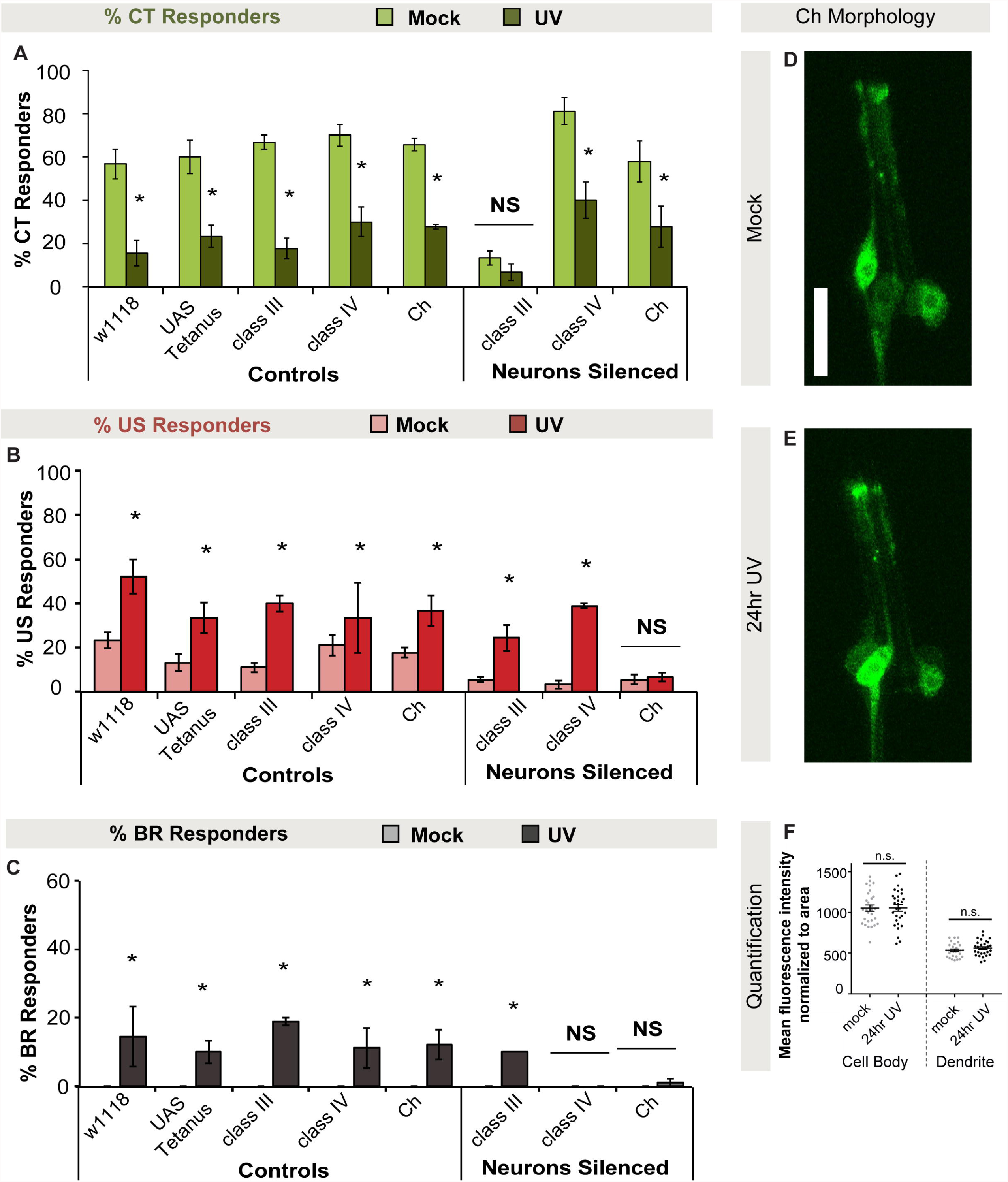
The UV-induced switch of cold-evoked behaviors requires class IV and Ch peripheral sensory neurons. (A-C) Larvae expressing an active or inactive (control) form of the tetanus toxin transgene (see materials and methods) in class III (CIII), class IV (CIV) or chordotonal neurons (Ch) were tested for cold-evoked behaviors 24 hours after UV exposure. Percent of CT (A), US (B), or BR (C) responders were averaged and UV- versus mock-treated responses were compared. n = 90. (D-E) Representative *in vivo* confocal microscopy images of mock treated larvae (D) and larvae 24 hours post UV treatment (E). Larval chordotonal morphology was visualized via *iav-Gal4>UASmCD8::GFP.* Scale bar: 20 μm. (F) Mean fluorescence intensity normalized to area for cell body and dendrites, where n = 29-30 neurons. Data are presented as the average ± s.e.m (A-C,F). Stats: (A-C) two-tailed Fisher’s Exact test * = p < 0.05. (F) Two-tailed t-test, n.s. = not significant, comparisons were between UV and mock control.

### UV irradiation does not alter cold-evoked calcium responses

To investigate whether there are any calcium changes at the sensory neuron level, we expressed GCaMP in relevant sensory neuron subtypes in mock and UV-irradiated third instar larvae and exposed these larvae to noxious cold. Consistent with previous findings [25], in mock treated larvae, CIII sensory neurons displayed a robust calcium response to noxious cold compared to baseline levels (Fig 4A, B, E). After UV irradiation, CIII sensory neurons still exhibited strong cold-evoked calcium responses, albeit with a slight, but non significant decrease in GCaMP response relative to mock CIII controls (Fig 4A, B, E). Interestingly, in mock-treated larvae, Ch neurons showed similar magnitude calcium responses to noxious cold as observed in CIII neurons (Fig 4A, C, F). As with CIII sensory neurons, Ch neurons still had robust cold-evoked calcium reponses after UV irradiation and a slight, but non significant, reduction in max ΔF compared to mock-treated Ch neurons (Fig 4A, C, F). Lastly, we investigated calcium responses in mock-treated CIV sensory neurons. Consistent with previous results [26], the CIV calcium response to noxious cold is significantly lower than in CIII and Ch sensory neurons (p<0.0001; two-way ANOVA with Sidak mulitple correction test). In UV-irradiated larvae, CIV sensory neurons have a slightly higher, but non significant, calcium response compared to mock treated neurons (Fig 4A, D, G). Together, these results suggest that the switches in noxious cold-evoked behavioral output following UV irradiation appear independent of alterations in calcium physiology in sensory neuron somata.

**Fig 4.**
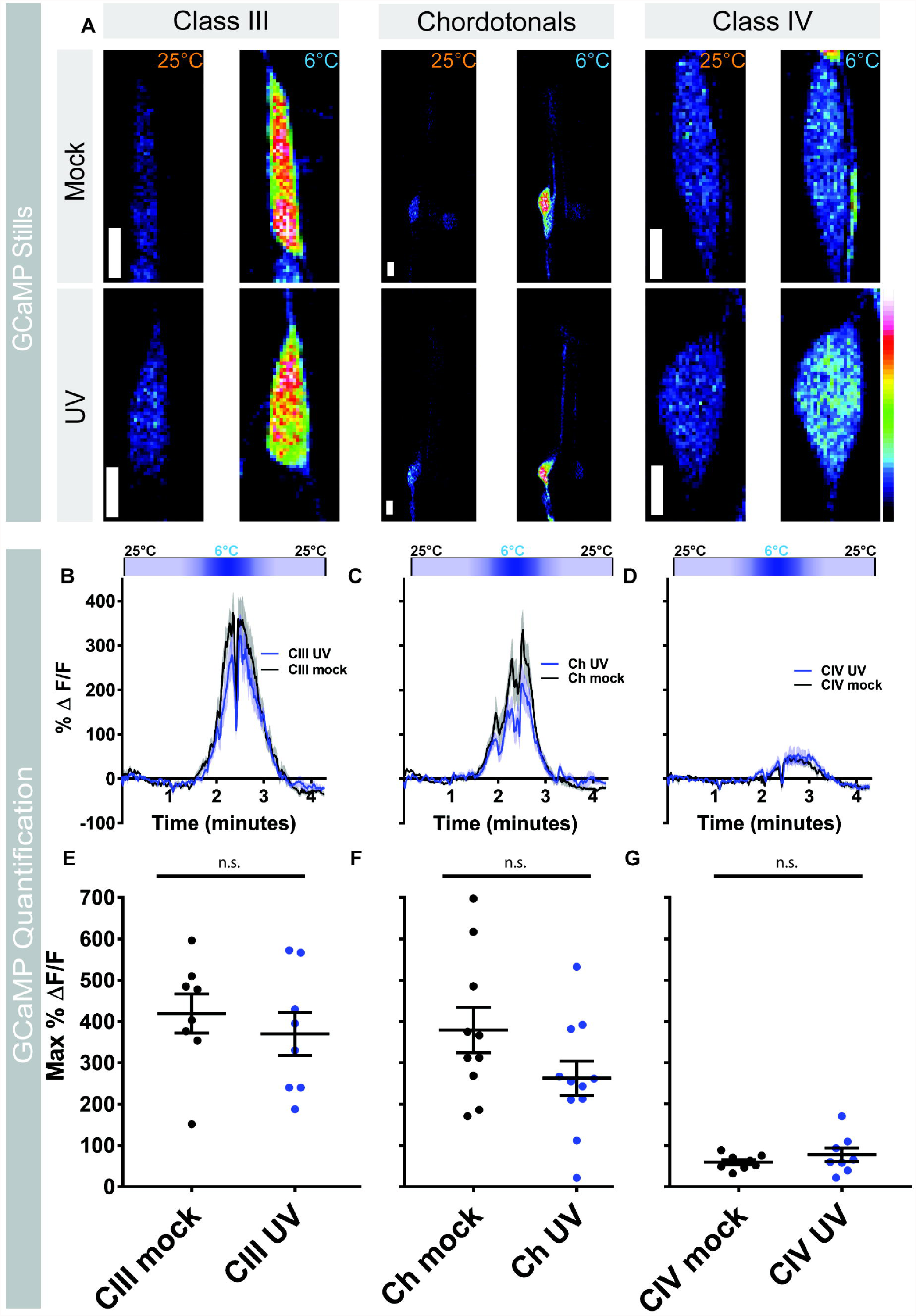
UV irradiation does not alter cold-evoked calcium responses. (A) *in vivo* confocal stills of larval CIII, Ch, and CIV sensory neurons expressing GCaMP6m at 25°C and 6°C in mock treated larvae and 24 hours post UV treatment. Scale bar: 5 μm. (B-D) Change in GCaMP6m fluorescence (ΔF) over time for CIII (B), Ch (C), and CIV sensory neurons (D). Coldmap on top of each graph represents stimulus temperature. Data represented as mean GCaMP6 response (black and blue trace lines) ± s.e.m (grey). (E-G) Max change in GCaMP6m fluorescence for mock- and UV-treated larvae 24 hours post-irradiation for CIII (E), Ch (F), and CIV sensory neurons (G), where the middle line is mean ± s.e.m. and n= 8-11 larvae. Stats: Two-tailed t-test (E-G), where the comparisons are between mock and UV treated conditions. n.s. = not significant.

### TRP channels are required for UV-induced changes in responses to cold in a class specific manner

TRP channels mediate a multitude of thermosensory responses in *Drosophila* and other animals [15]. We therefore hypothesized that changes in cold responses after UV may depend on TRP channels expressed in specific sensory neurons involved in sensitization to cold. TRP channels are expressed on class IV and chordotonal neurons [26,31,59] and function in these cells to mediate detection of sensory stimuli. In particular, we examined the TRPP channel genes *Pkd2* and *Brv1. Pkd2* has been shown to be required for baseline responses to cold and confers cold sensitivity to noncold-sensing neurons [26], while Brv channels have been implicated in cold avoidance in adult flies [30].

To test whether these TRPs are required for UV-induced changes in cold responses, we assayed mutants of *Pkd2* and *brv1* over relevant deficiencies for cold responses 24 hours after UV. Larvae mutant for *Pkd2* or *brv1* exhibited decreases in CT responders (Fig 5A), and increases in US responders (Fig 5B) to the cold probe after UV similar to mock controls. The mutant UV responses, however, are significantly altered when compared to irradiated control larvae for both CT (Fig 5A, red asterisks) and US responses (Fig 5B, red asterisks). Larvae mutant for *Pkd2* or *brv1* did not exhibit the increase in BR response after UV seen in wildtype controls (Fig 5C).

**Fig 5.**
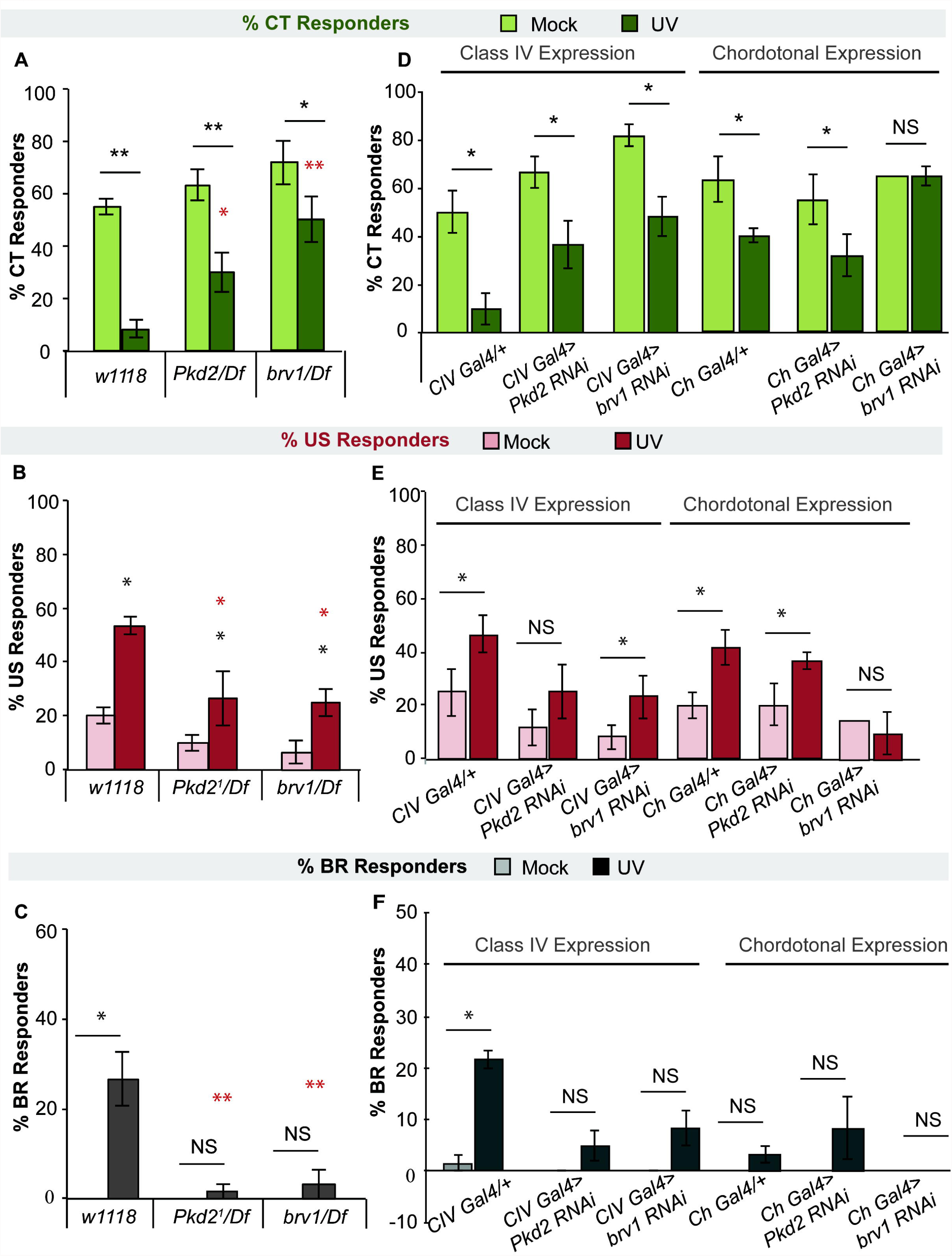
*Pkd2 and brv1* are involved in the UV-induced shift of behavioral responses to cold. (A-C) Percent of CT (A), US (B) or BR (C) responders to the cold probe (10 °C) 24 hours after UV in TRP channel mutants for *Brv1* or *Pkd2* over relevant deficiencies (Df). (D-F) Percent of CT (D), US (E), or BR (F) responders to the cold probe 24 hours after UV in larvae expressing *Pkd2* or *brv1-RNAi* transgenes in class IV or chordotonal neurons. (A-F) n = 3 sets of 20 larvae averaged ± s.e.m., except for (E) where *ChGal4>brv1-RNAi* was an n = 20-40 larvae. Stats: two-tailed Fisher’s exact test, * = p < 0.05, ** = p < 0.001. Black asterisks indicate statistical analysis between mock and UV groups for a given genotype. Red asterisks indicate statistical analysis between *w^1118^* UV-treated larvae and mutant UV-treated larvae. NS = no significance between UV and mock of same genotype.

To determine where these TRP channels may be required for cold sensitization, we examined larvae expressing gene-specific *UAS-RNAi* transgenes in CIV or Ch neurons. Interestingly, while *Pkd2^RNAi^* or *brv1^RNAi^* had no effect on the UV-induced decrease of CT responders when expressed in CIV neurons, this decrease was completely blocked when *brv1^RNAi^* was expressed in Ch neurons (Fig 5D). While irradiated larvae expressing *brv1^RNAi^* in CIV neurons still showed an increase in US responders compared to mock controls, *Pkd2^RNAi^* in CIV neurons attenuated this increase in US responders (Fig 5E). In the opposite manner, larvae expressing *brv1^RNAi^* in Ch neurons showed a blocked increase in US responders after UV compared to mock controls, while *Pkd2^RNAi^* expressing larvae exhibited a normal increase in US responders after UV (Fig 5E). For BR responses, larvae expressing *brv1^RNAi^* or *Pkd2^RNAi^* in CIV neurons also had blocked increases in BR responses after UV compared to mock controls (Fig 5F). It is more difficult to discern the contributions of Pkd2 and Brv1 in Ch neurons however, since although no increase in BR responses was observed upon expression of *Pkd2^RNAi^* or *brv1^RNAi^*, it was also quite low in the Gal4 alone control (Fig 5F). Collectively, these data suggest that Pkd2 is important in CIV for both increased US and the emergence of BR responses after UV, while Brv1 acts in CIV and Ch neurons to mediate increases in US responses and the decreases in CT responses to cold after UV.

## Discussion

We show that after UV-induced epidermal tissue damage *Drosophila* larvae exhibit altered responses to cold stimuli. They do so in a complex manner involving a shift in behavioral output away from CT and towards US and BR responses. The increase in US responses appears to require Ch neurons while the emergence of BR responses to cold requires both CIV and Ch neurons (see summary, Fig 6). Molecularly, the TRPP type channels, Pkd2 and Brv1, are required for the observed UV-induced behavioral shifts in response to noxious cold acting in CIV and/or Ch neurons.

**Fig 6.**
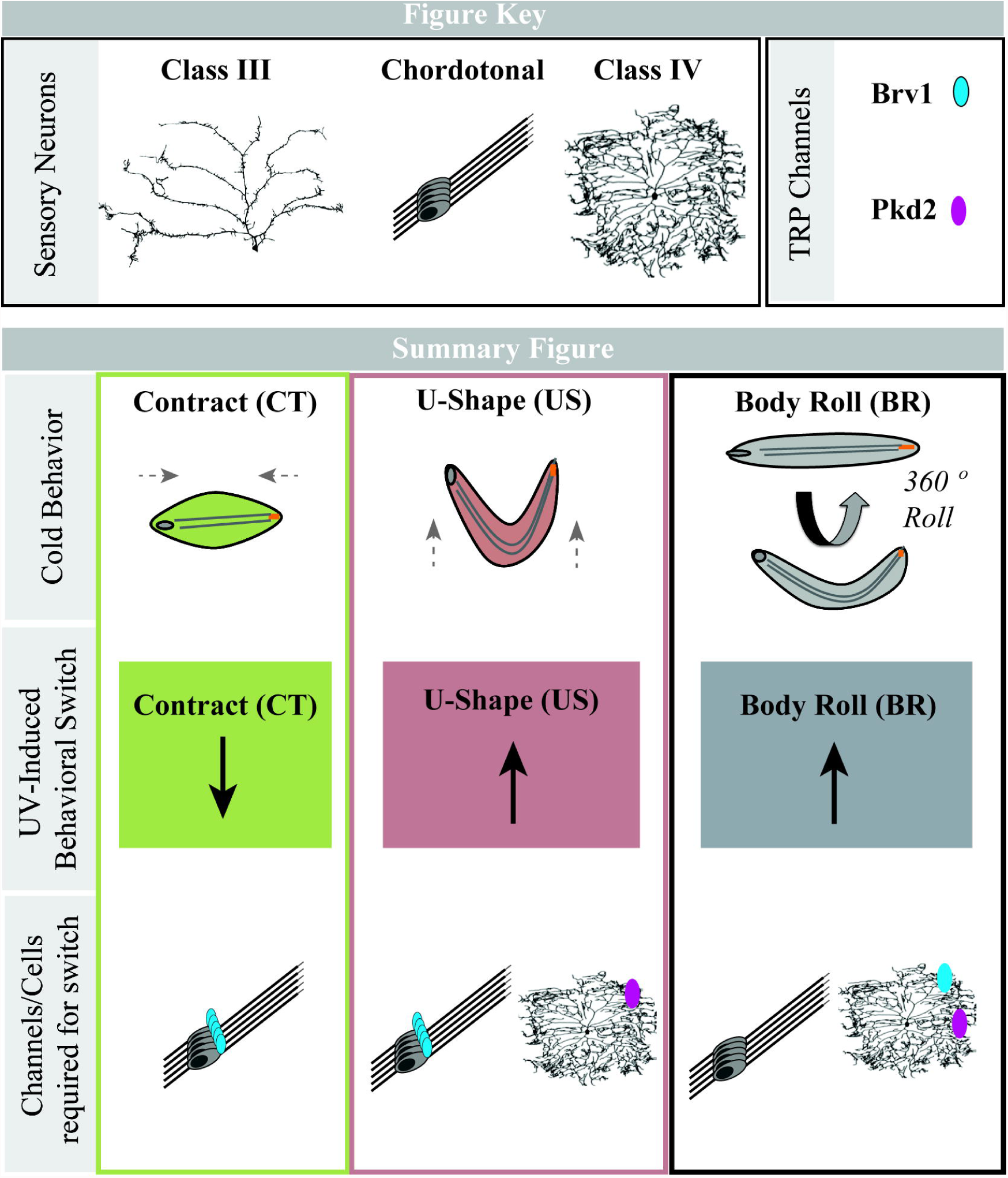
Summary Figure of sensory neurons and genes required for UV-induced behavioral shift(s) in cold-evoked behaviors in *Drosophila* larvae. The top two panels provide a key, marking indicators for the specific cell types (Class III, Chordotonal and Class IV) and TRP channels (Brv1 in blue, Pkd2 in purple) used in the lower panels. The first row illustrates the baseline cold-evoked behaviors: a full body contraction (CT), a raising of the head and tail into a “U-Shape” (US), and a 360° lateral body roll (BR). The middle panel indicates the shift in each of these behaviors observed in larvae 16-24 hours after being exposed to UV. The last panel indicates the TRP channels and sensory neuron types involved in each of these UV-induced behavioral shifts.

Morphological alterations following UV-induced injury could potentially contribute to cold sensitization behavioral shifts. However, we demonstrate that neither CIII nor Ch neurons exhibit morphological changes when comparing mock- and UV-irradiated larvae. Previous studies using a similar irradiation paradigm reveal no UV-induced change in CIV dendritic morphology [46].

Physiologically, cold-evoked calcium analysis in mock versus UV treated larvae revealed that the UV-induced behavioral switch appears to occur independently of calcium dynamics in CIII, Ch or CIV sensory neurons. These analyses did, however, reveal that Ch neurons, like CIII neurons, robustly respond to noxious cold under native conditions (in the absence of injury). As we did not observe any significant changes in mean calcium dynamics with UV treatment at the level of the primary sensory neurons, the observed shifts in behavioral output in response to noxious cold following injury may be an effect mediated at the synapse or downstream circuit(s). For example, previous studies have demonstrated that co-activation of Ch neurons and CIV neurons via vibration and noxious heat, respectively, synergistically increases the BR response, suggestive of an interaction between these sensory circuits [63]. Moreover, CIII and CIV axons terminate in adjacent regions of the neuropil [64] and recent studies have documented some common postsynaptic first order interneuron targets shared by CIII and CIV neurons that can serve as integrators in promoting specific somatosensory behavioral responses [65,66]. UV-induced injury may lead to complex changes in neural circuit processing of the noxious cold stimulus where use of alternate neurotransmitters or changes in the quantal release of neurotransmitters from sensory neurons could possibly lead to a behavioral switch. Another possibility, less likely, could be a weakening of CIII synapses to yet unknown first order interneurons, thus leading to alternate behaviors. Finally, UV exposure could lead to alterations in downstream postsynaptic interneuron targets that may serve as integrators of multiple sensory neuron subtypes or perhaps affect motor neuron activity that may indirectly affect muscle contractility.

Pkd2, recently discovered to be required for baseline CT responses to noxious cold [26], here appears to also act in CIV for UV-induced shifts in US and BR responses to cold. Interestingly, *brv1* functions in CIV neurons to mediate the emergence of BR responses, while in Ch neurons Brv1 appears to mediate the observed decrease in CT responders and increase in US responses to cold after UV damage. The fact that we see an overall decrease in cold-evoked CT in the irradiated *brv1* mutant, yet when *brv1-RNAi* is expressed in Ch neurons this decrease is blocked, suggests that perhaps opposing roles of *brv1* in different neuronal subtypes balances the ultimate behavioral response produced. This data indicates that both of these TRP channels play a role in observed behavioral shifts from CT responses to US and BR responses to cold after UV, in two distinct populations of peripheral sensory neurons.

Why does this shift from the dominant cold response (CT) towards two alternative behaviors (US and BR) following UV-induced tissue damage occur? Given that UV exposure causes epidermal damage and apoptosis [38], and that the CT response requires a significant change in body length [26], it may be that a full-body contraction is physically more “painful” than a US or BR. Further, it may be that a CT response exacerbates epidermal damage more than US or BR, making it advantageous for the larva to avoid CT in favor of other cold-evoked behaviors. We have previously shown that thermogenetic activation of CIII neurons, via expression of TRPA1 and application of heat (45 °C), and CIV neurons (by the heat alone), results in an approximate balance of CT and BR responses [26], suggesting that crosstalk between these neurons may be present.

Mechanistically, the behavioral shift from CT to US and BR could result from UV altering the sensitivity, localization, and/or expression level of Pkd2 and/or Brv1 in CIV and/or Ch neurons. This is particularly interesting for CIV neurons, since ectoptic expression of *Pkd2* in these normally cold-insensitive neurons confers cold sensitivity [26]. While many TRP channels are expressed on CIII neurons [26,59], specific changes in TRP channel sensitivity or expression could presumably alter neuronal function or result in a suppression of activity, allowing the CIV neuron behavior (BR) to be observed at the expense of other behaviors coming from other (CIII and Ch) neurons. For Ch neurons, it remains unclear which channels may be required for the observed emergence of BR responses to cold, but only *brv1* appears to be required for US responses as well as the decrease in CT responses after UV. Currently, it is unknown whether these genes, particularly *brv1,* are expressed in Ch neurons.

Together, we illustrate an intriguing shift in cold-evoked responses following UV-induced tissue damage. This shift involves, at least in part, CIV sensory neurons, similar to the sensitization of BR responses to noxious heat after UV-induced tissue damage [38,44]. However, there is an interesting dichotomy in the genes required for cold-induced sensitization that appears different from those that mediate sensitization to noxious heat. In the case of UV-induced sensitization to noxious heat, TRPA type channels are involved [44], whereas here we show that TRPP type channels, *Pkd2* and *brv1,* are required for UV-induced shifts in behavioral responses to noxious cold. In all, this work establishes that *Drosophila* can be used to study nociceptive cold sensitization and to identify key players in the process. The emergence of BR responses to cold we see here may be an interesting model for advancing studies where cold is perceived as a “burning” sensation [67]. In the thermal grill illusion, a subject is instructed to place their hand on a grill made up of interlaced warm (40 °C) and cool (20 °C) bars [68]. Instead of perceiving either warm or cool individually or simultaneously, the illusion instead leads to the surprising perception of noxious cold as a burning sensation [68]. It is thought that the study of this illusion can yield important insights into pain management and relief [69] because it illustrates, like our data here, that the line between the perception of temperature versus pain can be shifted under different environmental contexts, indicating that the circuits for these pathways are likely to overlap. Indeed, when CIII and CIV neurons are optogenetically activated simultaneously, the resulting behavioral response is a CT, revealing that a circuit level shift in behavior is possible under certain circumstances.

There is a large clinical need for a better understanding of the mechanisms that underlie sensitivitiy to cool and noxious cold stimuli, particularly how the perception of cool stimuli can be sensitized following injury or in models of chronic neuropathic pain. Utilizing *Drosophila,* and the tool and assay described here, will aid research in identifying novel genetic players and mechanisms underlying cold nociception and nociceptive sensitization to better inform our understanding of pain syndromes in the future. Together this study creates a novel platform to study nociceptive cold sensitization in *Drosophila* larvae and reveals an interesting shift in behavioral output to cold stimuli upon tissue injury that requries specific sensory neurons and TRP channel genes. Further investigation into the mechanisms of cold sensitization using these tools and assays will undoubtably aid our understanding of the mediators that may be involved in clinical cases of cold hypersensitization.

## Contributions

Conceptualization: HNT, AAP, DNC, MJG; Methodology: HNT, AAP, DNC, MJG; Formal analysis: HNT, AAP; Investigation: HNT, AAP; Resources: DNC, MJG; Writing – original draft: HNT, MJG; Writing – review & editing: HNT, AAP, DNC, MJG; Visualizations: HNT, AAP; Supervision: DNC, MJG; Project administration: DNC, MJG; Funding acquisition: HNT, AAP, DNC, MJG.

## Acknowledgements

We acknowledge the VDRC stock center for *UAS-RNAi* transgenes and Marco Gallio, Teri Watnick, Kartik Venkatachalam, and Yuh Nung Jan for fly stocks. Stocks obtained from the Bloomington *Drosophila* stock center (NIH P40ODO18537) were used in this study. We also thank members of the Cox and Galko labs for critical evaluation of the manuscript. The authors declare no competing financial interests. This research was supported by the National Institute of Neurological Disorders and Stroke (NINDS) (R01 NS086082 to D.N.C. and the National Institute of General Medical Sciences (NIGMS) (R35 GM126929 to M.J.G.); an NIH Predoctoral Kirschstein National Research Service Award (NRSA) Fellowship (NINDS F31 NS083306) and Marilyn and Frederick R. Lummis, Jr. MD Fellowship (H.N.T.); Georgia State University (GSU) 2CI Neurogenomics Fellowship and Kenneth W. and Georgeanne F. Honeycutt Fellowship to A.A.P; and a GSU Brains and Behavior grant to D.N.C.

